# The pharmacological perturbation of brain zinc impairs BDNF-related signalling and the cognitive performances of young mice

**DOI:** 10.1101/267609

**Authors:** Valerio Frazzini, Alberto Granzotto, Manuela Bomba, Noemi Massetti, Vanessa Castelli, Marco d’Aurora, Miriam Punzi, Mariangela Iorio, Alessandra Mosca, Stefano Delli Pizzi, Valentina Gatta, Annamaria Cimini, Stefano L. Sensi

## Abstract

Zinc (Zn^2+^) is a pleiotropic modulator of the neuronal and brain activity. The disruption of intraneuronal Zn^2+^ levels triggers neurotoxic processes and affects neuronal functioning. In this study, we investigated how the pharmacological modulation of brain Zn^2+^ affects synaptic plasticity and cognition in wild-type mice. To manipulate brain Zn^2+^ levels, we employed the Zn^2+^ (and copper) chelator 5-chloro-7-iodo-8-hydroxyquinoline (clioquinol, CQ). CQ was administered for two weeks to 2.5-month-old (m.o.) mice, and effects studied on BDNF-related signalling, metalloproteinase activity as well as learning and memory performances. CQ treatment was found to negatively affect short- and long-term memory performances. The CQ-driven perturbation of brain Zn^2+^ was found to reduce levels of BDNF, synaptic plasticity-related proteins and dendritic spine density *in vivo*.

Our study highlights the importance of choosing “when”, “where”, and “how much” in the modulation of brain Zn^2+^ levels. Our findings confirm the importance of targeting Zn^2+^ as a therapeutic approach against neurodegenerative conditions but, at the same time, underscore the potential drawbacks of reducing brain Zn^2+^ availability upon the early stages of development.

## Introduction

The neurotrophin brain-derived neurotrophic factor (BDNF) critically modulates the plasticity of the Central Nervous System (CNS) with a broad spectrum of activities that spread from neuronal differentiation and survival to synaptogenesis and activity-dependent synaptic plasticity ^1,2^.

BDNF is synthesized as pre-pro-BDNF, the precursor form, cleaved into pro-BDNF ^3–5^, released in the extracellular space, and then cleaved into its mature counterpart (mBDNF) by extracellular proteases like plasmin or metalloproteinases (MMPs), like the MMP-2 and 9 ^6–9^. mBDNF and its preferred receptor TrkB play a crucial role in the plasticity-related morphological modifications of spines. Regulation of spine morphology represents the physical basis of learning and memory processes, a phenomenon defined as structural plasticity. The balance between pro-BDNF and mBDNF has significant biological implications as the immature and mature forms of the protein, by activating distinct class of receptors ^10,11^, exert different actions on synaptic plasticity and neuronal survival ^12,13^. mBDNF promotes neuroprotective activity in *in vitro* and *in vivo* preclinical models of neurodegeneration ^14–16^. These preclinical findings are paralleled by evidence in human studies that indicate that high serum levels of mBDNF, as well as high *BDNF* gene expression, are associated with a reduced risk of dementia in elderly subjects and a milder disease course in dementia patients ^17–19^.

The metalloproteinase, MMP-9, is highly enriched in the hippocampus, the cerebral cortex, and the cerebellum ^20^, brain regions where the enzyme is localized in the extracellular matrix near the dendritic spines of excitatory synapses ^21^. MMP-9 contributes to learning, memory and neuronal plasticity by promoting the spine remodelling and immobilization of ionotropic receptors that occur upon long-term potentiation (LTP) stimulation ^22,23^. In preclinical models of Alzheimer’s disease (AD), the functional overexpression of MMP-9 increases the brain levels of mBDNF and prevents the development of AD-related cognitive deficits ^24,25^. MMP-9 is structurally characterized by a 3-histidine zinc-binding catalytic motif (His-Glu-Xaa-Xaa-His) and requires zinc (Zn^2+^) to exert its proteolytic activity ^26^.

Zn^2+^ is largely present in the CNS, stored within synaptic vesicles at several glutamatergic nerve terminals, and synaptically released upon neuronal activity ^27,28^. Zn^2+^ affects neuronal processes as well as BDNF signalling ^29^. The metal, through MMP activation, promotes the cleavage of pro-BDNF to mBDNF ^8,9^. Zn^2+^ also modulates BDNF signalling without affecting the MMP activity. This activity is achieved by direct transactivation of TrkB, the preferred and high-affinity BDNF receptor ^30^. In addition, *in vivo* administration of Zn^2+^ increases the cortical levels of the BDNF mRNA ^31^, thereby indicating a pro-transcriptional activity of the cation.

In a preclinical transgenic AD model, we have previously shown that long-term dietary supplementation of Zn^2+^ promotes the activation of MMP-2 and MMP-9, increases the brain levels of mBDNF, and counteracts the development of AD-related pathology and memory deficits ^32^.

In this study, we investigated how the pharmacological modulation of brain Zn^2+^ affects BDNF signalling, synaptic plasticity, and cognition in young mice. To manipulate brain Zn^2+^ levels, we employed the Zn^2+^ (and copper) chelator 5-chloro-7-iodo-8-hydroxyquinoline (clioquinol, CQ) as previous studies have shown that the intraperitoneal administration of CQ promotes a rapid reduction in the brain pool of chelatable Zn^2+ 33–38^. CQ was administered for two weeks to 2.5-month-old (m.o.) mice and effects studied on BDNF-related signalling, MMP activity as well as learning and memory performances.

## Results

### CQ administration impairs learning, short- and long-term memory performances in young mice

Male C57Bl/6 mice were treated, for up to 2 weeks, with daily intraperitoneal (i.p.) injections of CQ (30 mg/kg). The rationale for choosing this concentration and route of administration was threefold: 1) this CQ concentration has been shown to be devoid of significant side effects in rodents including changes in body weight (Fig. 1B; 25.8±1.6 g in control vs 24.9±0.91 in CQ treated mice, p=0.64); 2) the dose matches the one employed in clinical trials; 3) the i.p. route is highly effective in reducing the pool of chelatable Zn^2+^ in the brain and the hippocampus in particular ^33,39^. In agreement with previous studies ^33–38^, we obtained a significant decrease of hippocampal Zn^2+^ levels (Supplementary fig. 1; normalized TSQ fluorescence in control: 1.00±0.08 vs 0.49±0.04 in CQ-treated mice, p<0.001).

**Figure 1.**
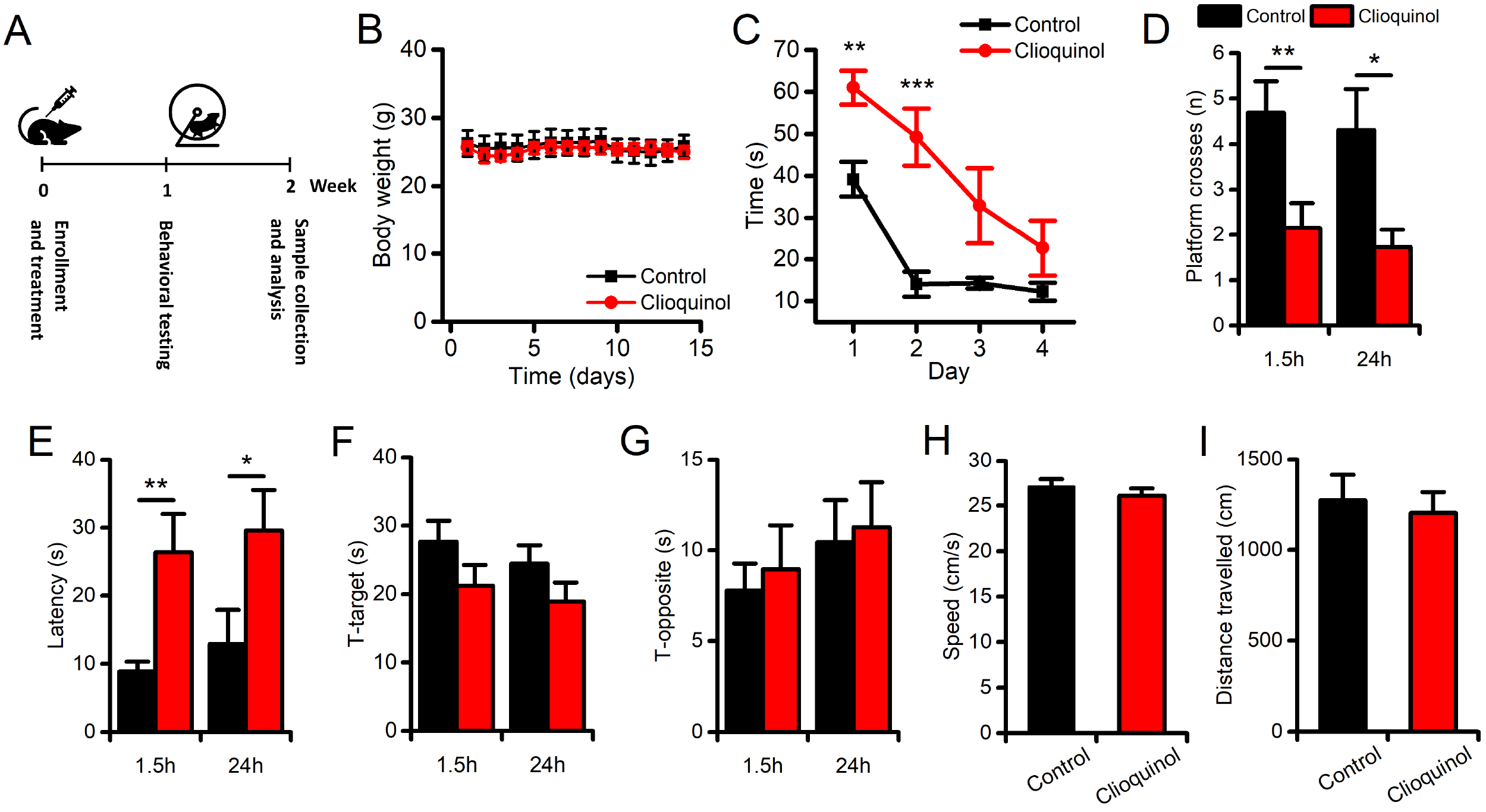
CQ administration impairs learning and memory performances in mice. Spatial memory performances were evaluated with the MWM test. (A) The pictogram illustrates the study design. (B) The graph illustrates the daily body weight monitoring of vehicle- and CQ-treated mice during the 2-week treatment period (n=8 per study group). (C) The graph illustrates the learning curve of the vehicle- and CQ-treated mice (n=9 and n=10, respectively) as evaluated during the 4-day training session. Compared to vehicle-treated animals, CQ-treated mice showed significantly impaired learning performances in the first 3 trials of the training sessions. Note that the impairment disappeared in the fourth day of training. (D) The bar graph shows CQ-driven decreases in the number of platform crosses (the number of times the mouse crosses the location where the platform used to be) in the 1.5h (to evaluate STM) and the 24h (to evaluate LTM) trials. (E) The bar graph shows CQ-driven increases of latency (the time spent to reach the location where the platform used to be) in both the STM and the LTM trials. (F-G) Bar graphs show the absence of drug-related changes in the time spent in the target (the quadrant where the platform used to be) or the opposite (the quadrant opposed to the one where the platform used to be) quadrants, in the STM and LTM trials (F and G, respectively). (H) The bar graph shows no CQ-driven changes in swimming speed in LTM trials in a subset of mice (n=5 for control and n=4 for CQ-treated mice). (I) The bar graph shows no CQ-driven changes in distance travelled in LTM trials in a subset of mice (n=5 for control and n=4 for CQ-treated mice). Data are presented as mean ± standard error of the mean (SEM). “*” indicates p < 0.05, “**” indicates p < 0.01.

The effects of CQ or vehicle on hippocampus-dependent spatial learning were assessed, at the end of the second week of treatment, with the Morris Water Maze (MWM) test, a widely employed task that evaluates memory performances in rodents ^40^ (Fig. 1A). Compared to vehicle-treated mice, CQ-treated animals displayed learning impairment. CQ-treated mice showed a statistically-significant increase in the time spent to reach the platform upon the first two training sessions (Fig. 1C; day 1 p<0.01, day 2 p<0.001). These learning deficits were transient and disappeared in the final two days of training (Fig. 1C; day 3 p=0.069, day 4 p=0.16). The analysis of memory performances revealed that, compared to vehicle-treated animals, CQ-treated mice performed significantly worse in Short- (1.5 h) and Long-Term (24 h) Memory (STM and LTM, respectively) skills. The CQ-treated group showed a decreased number of platform crosses (Fig. 1D; for STM: 4.69±0.69 in control vs 2.15±0.54 in CQ-treated mice, p=0.008; for LTM: 4.3±0.9 in control vs 1.73±0.38 in CQ-treated mice, p=0.04) and longer times to reach the platform location (Fig. 1E, escape latency for STM: 8.84±1.5 s in control vs 26.38±5.6 s in CQ-treated mice, p=0.007; escape latency for LTM: 12.9±5 s in control vs 29.58±5.9 s in CQ-treated mice, p=0.01). No significant differences, between the two study groups, were observed as far as the time spent in the target or opposite quadrants (Fig. 1F-G, T-target, and T-opposite, respectively). Additional analysis showed no significant differences between the two study groups regarding swimming speed or the total distance travelled upon the LTM probe trial (Fig. 1H-I; p=0.43 for the swimming speed and p=0.71 for the distance travelled), thereby indicating that the CQ-mediated effects were not driven by decreased motivational cues or impaired swimming abilities.

### CQ reduces BDNF signalling in treated mice

Zn^2+^ has been shown to promote pro-BDNF maturation *in vitro* ^8^. Western blot (WB) analysis of brain samples revealed a significant decrease of BDNF levels in CQ-treated mice when compared to vehicle-treated animals (Fig. 2B; see Supplementary Table 1 for the statistics of the WB data). The reduction is region-specific and occurred in the hippocampus, cortex, and striatum but not in the cerebellum. These regional changes were matched by parallel decreased levels of the preferred BDNF receptor, TrkB (Fig. 2C). To assess whether BDNF and TrkB reductions affect BDNF-related signalling, we measured brain levels of the activated-phosphorylated-form of TrkB (pTrkB) and ERK5 (pERK5), a downstream effector of the BDNF-TrkB signalling cascade ^41^. WB analysis showed no significant differences between CQ- and vehicle-treated animals in the pTrkB/TrkB ratio (Fig. 2D) while CQ administration induced a significant decrease in pERK5 levels (Fig. 2F).

**Figure 2.**
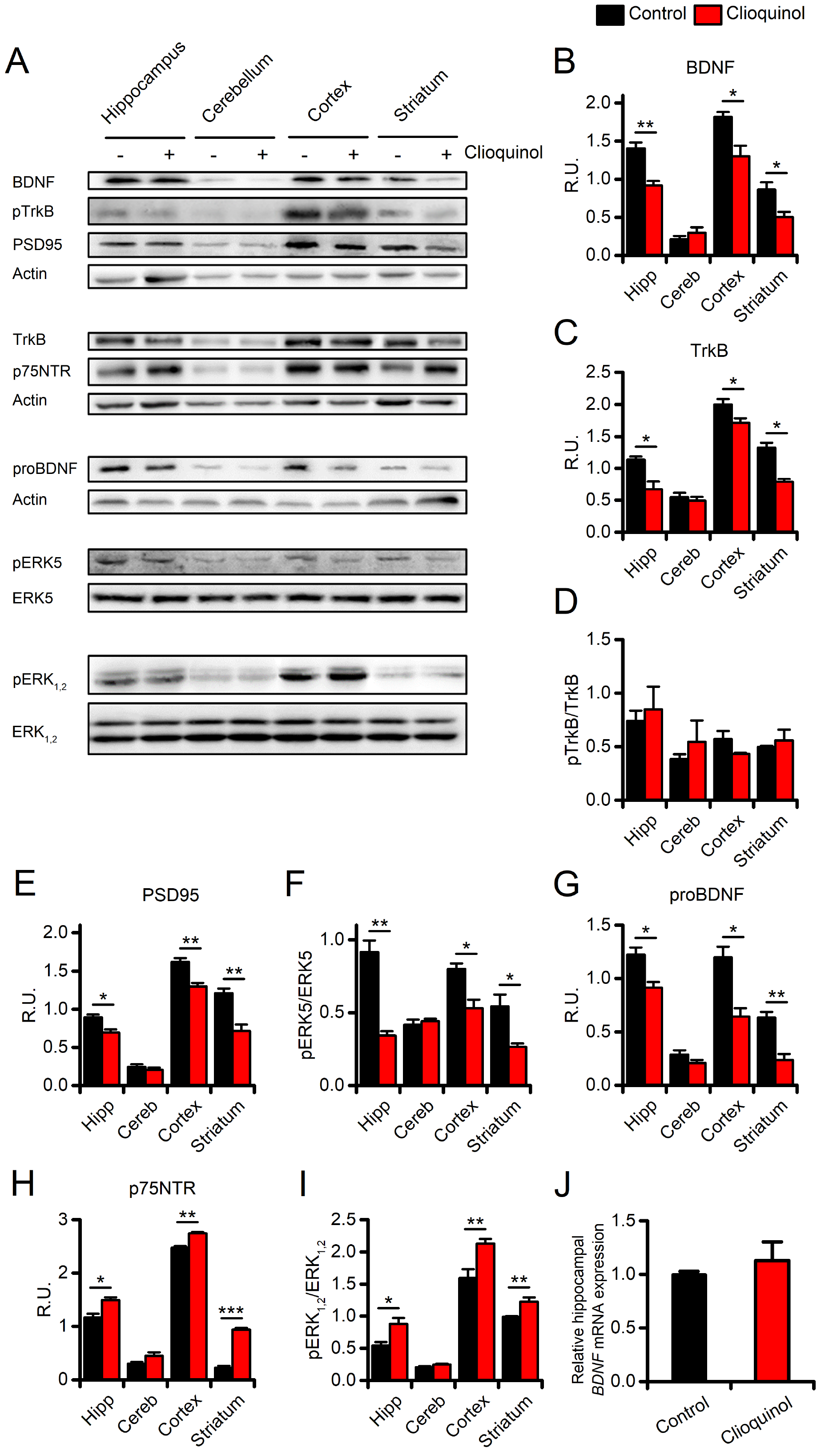
Effects of CQ administration on BDNF signalling. (A) Representative images of WB experiments showing vehicle- or CQ-driven effects on BDNF signalling in the hippocampus (Hipp), the cerebellum (Cereb), the cerebral cortex (Cortex), and the striatum of the two study groups. (B, C) Bar graphs show quantification of CQ-driven changes in mBDNF and TrkB expression levels. (D) Bar graphs show no drug-related changes in levels of TrkB phosphorylation (pTrkB/TrkB ratio) in the two study groups. (E) Bar graphs depict CQ-driven changes in PSD95 expression levels in the study groups. (F) Bar graphs depict levels of pERK5 in the study groups. (G, H) Bar graphs depict CQ-driven changes in proBDNF and p75NTR protein levels. (I) Bar graphs depict levels of pERK_1,2_ in the 2 study groups. All experiments were performed, at least, three times from independent samples. (J) The bar graph depicts mRNA levels of BDNF measured by real-time PCR in the vehicle- and CQ-treated mice (n=3 per condition). Data are presented as mean ± SEM. “*” indicates p < 0.05, “**” indicates p < 0.01, and “***” indicates p < 0.001.

The neurotrophin BDNF, by activating mechanisms related to functional and structural plasticity, is a potent modulator of LTM ^42^. We, therefore, investigated whether the CQ-driven reduction of BDNF and TrkB translates into changes in PSD95 levels, a marker of the post-synaptic integrity. Matching the CQ-related regional changes in BDNF, we found region-specific decreases of PSD95 in the hippocampus, cortex, striatum, but not in the cerebellum (Fig. 2E).

### CQ promotes p75NTR signalling in treated mice

Since pro-BDNF-p75NTR activation can modulate neurotoxic activities, we investigated whether the observed CQ-driven impairments of the BDNF-TrkB signalling axis are associated with changes in the pro-BDNF-p75NTR signalling cascade. WB analysis of these pathways showed that, compared to vehicle, CQ treatment promotes a region-specific increase of p75NTR levels along with high levels of phosphorylated ERK_1,2_ (pERK_1,2_; Figs. 2H-I). Surprisingly, the CQ-treatment was also found to decrease pro-BDNF levels in the same brain regions (Fig. 2G). The CQ-driven activation of the p75NTR signalling cascade did not produce overt signs of neurotoxicity *in vitro* and *in vivo* (Supplementary fig. 2).

### CQ treatment does not affect the hippocampal BDNF mRNA levels

Previous *in vivo* studies have shown that Zn^2+^ supplementation increases the mRNA expression of BDNF ^31^. On the contrary, pharmacological manipulations aimed at reducing the brain Zn^2+^ levels negatively modulate the BDNF mRNA levels ^43^. To address whether CQ-driven Zn^2+^ chelation interferes with the *BDNF* expression, we evaluated mRNA levels in the hippocampi of CQ- and vehicle-treated mice. Quantitative Real-Time PCR (qRT-PCR) analysis showed that CQ administration did not affect the hippocampal *BDNF* gene expression (Fig. 2J; 1.00±0.02 in control vs 1.12±0.17 in CQ treated mice, p=0.54).

### CQ treatment reduces dendritic spine density in vitro and ex vivo

As BDNF potently modulates synaptic plasticity ^30,44^, we first assessed, in an *in vitro* setting, whether CQ treatment alters the number of dendritic spines and/or their morphology. To that aim, EGFP-transfected primary hippocampal neurons were exposed to CQ (10 μM for 3 days) or vehicle and spines evaluated with high-resolution confocal microscopy. Confocal imaging analysis showed that, compared to vehicle-treated sister cultures, CQ-treated neurons underwent a significant reduction in dendritic spine density (Figs. 3A-B; spines per μm: 0.51±0.07 in control vs 0.22±0.02 in CQ-treated cultures, p=0.020). No differences were found, between the two study groups, regarding spine volumes, areas, diameters or lengths (Figs. 3C-F). To assess whether CQ also impairs spine pruning *in vivo,* we performed Golgi staining on brain slices of treated animals. Compared to vehicle-treated mice, the hippocampal CA1 pyramidal neurons of CQ-treated mice showed a significant decrease (≈ −30%) in their dendritic spine density (Fig. 3G-H; spines per μm: 1.41±0.13 in control vs 0.96±0.1 in CQ-treated mice, p=0.029).

**Figure 3.**
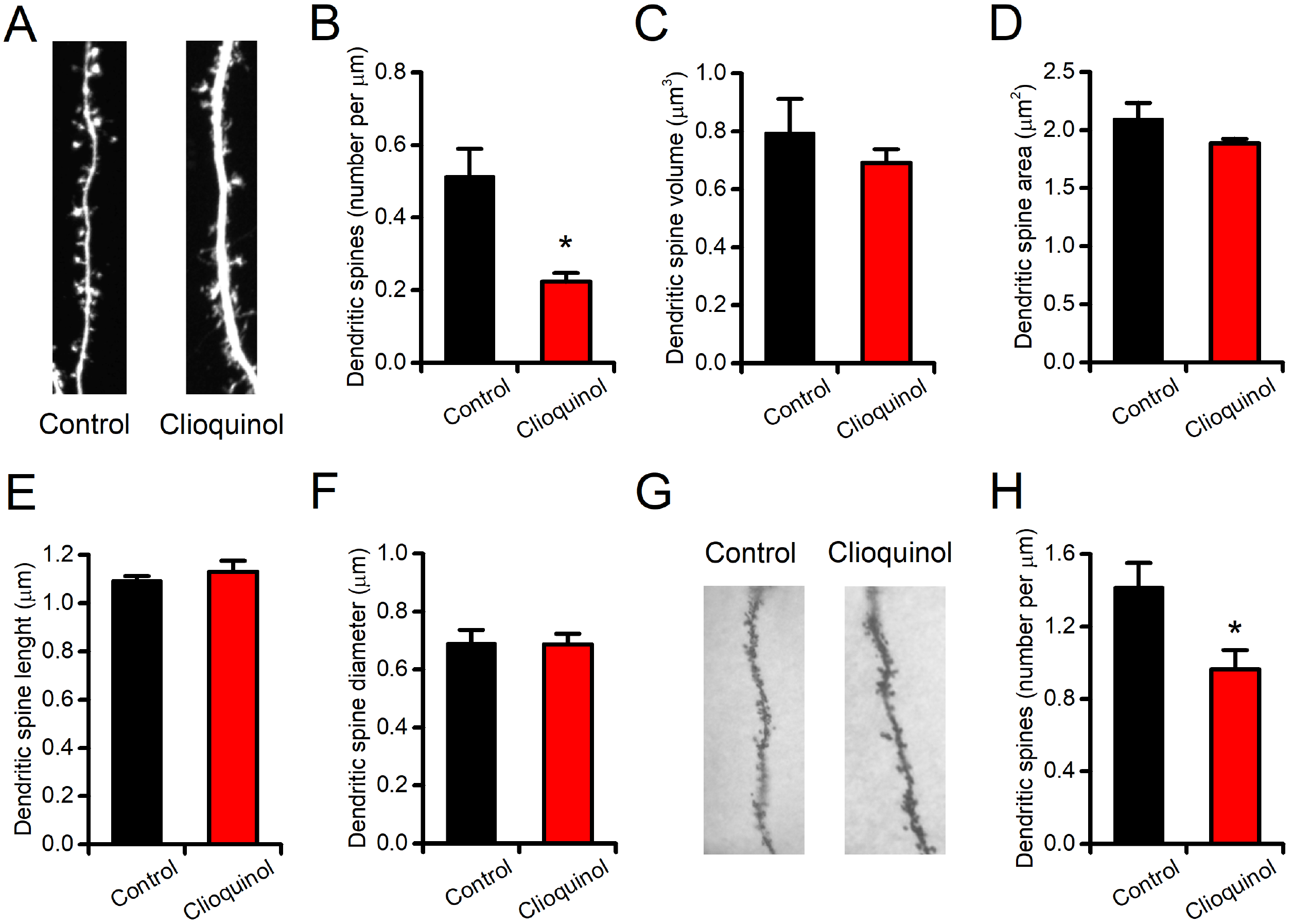
CQ reduces dendritic spine density *in vitro* and in *ex vivo* brain samples. (A) Representative images of proximal dendrites obtained from EGFP-transfected primary hippocampal neurons treated with vehicle (left) or CQ (10 μM for 3 days; right). (B) Bar graph depicts the quantification of dendritic spine density observed with confocal microscopy after the 3-day treatment period (n=4 for control and n=3 for CQ-treated neurons). (C-F) Bar graphs depict the quantification of additional parameters related to dendritic spine morphology as spine volumes (C), spine areas (D), spine lengths (E), and spine diameters (F). (G) Representative images of Golgi-stained CA1 pyramidal dendrites obtained from vehicle- and CQ-treated mice. (H) Bar graph depicts the quantification of dendritic spine density as observed at the end of the 2-week treatment period (n=19-35 neurons from 3 mice per condition). Data are presented as mean ± SEM. “*” indicates p<0.05.

### CQ reduces MMP activity in vitro

MMPs, and MMP-9 in particular, promote BDNF maturation through the proteolytic cleavage of pro-BDNF ^8^. To address whether CQ interferes with MMP activity we performed a gelatin zymographic essay on MMPs isolated from the medium of cultured hippocampal neurons. After gel electrophoresis, the MMP developing buffer was supplemented with CQ (10 μM) or the high-affinity Zn^2+^ chelator TPEN (100 μM, here employed as a positive control) and results compared with those of control gels. CQ almost completely abolished the MMP-9 and MMP-2 activity (Fig. 4; normalized MMP-9 activity: 1.00±0.09 in control vs 0.08±0.02 for CQ-treated gels, p<0.001; normalized MMP-2 activity: 1.00±0.17 in control vs 0.02±0.01 for CQ-treated gels, p<0.001). Gel treatment with TPEN did not induce further MMP reduction, thereby supporting the notion of a maximal inhibitory activity exerted by CQ (Fig. 4A-B).

**Figure 4.**
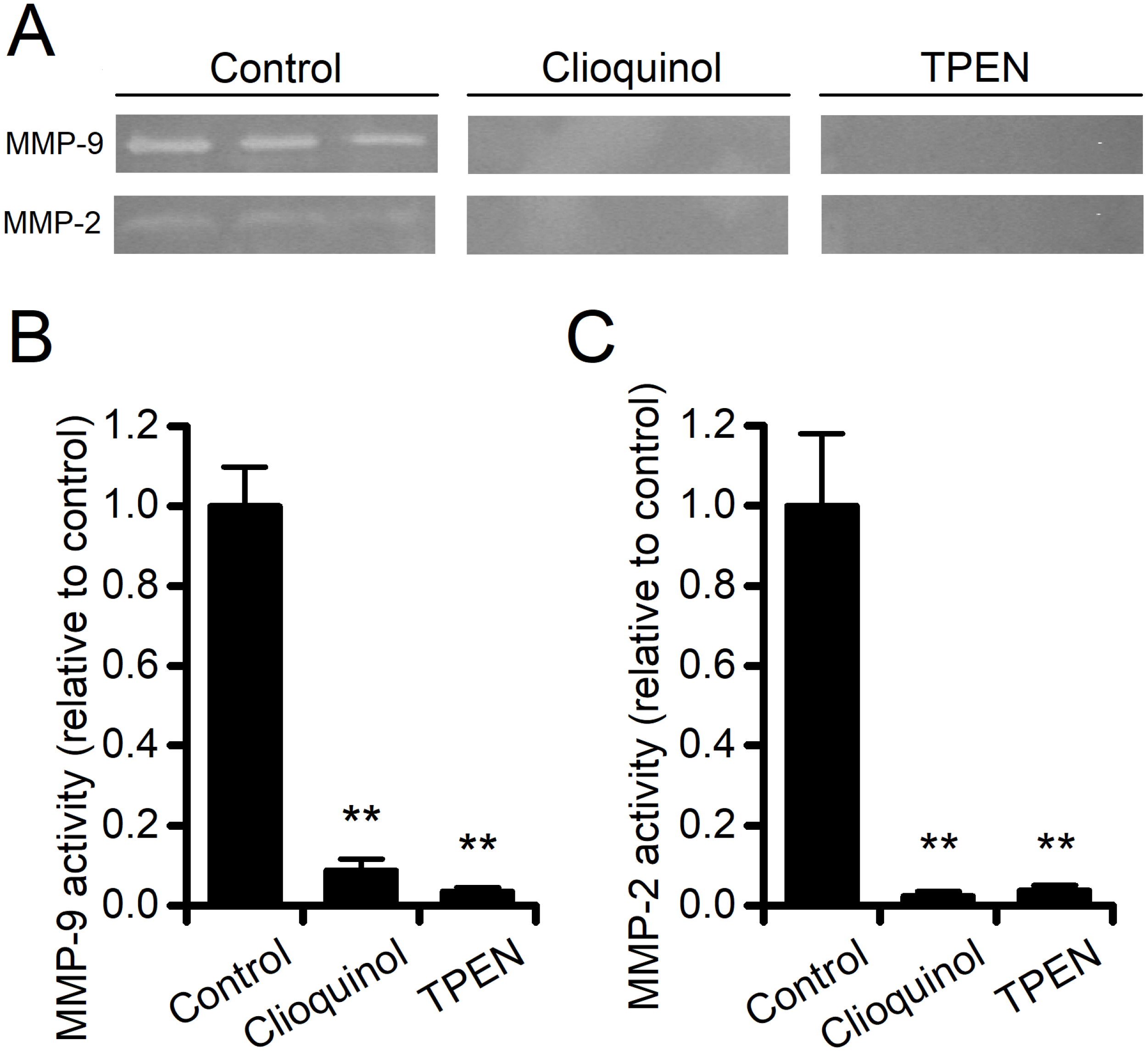
CQ reduces MMPs activity *in vitro*. (A) Representative images of gelatin zymography showing a CQ-driven decrease of MMP-9 and MMP-2 activity. Note that a CQ concentration as low as 10 μM has the same effect of TPEN (100 μM), here employed as a control. (B, C) Bar graphs show the quantification of images in A for MMP-9 and MMP-2, respectively (n=6 per condition). Data are presented as mean ± SEM. “**” indicates p<0.01.

## Discussion

In this study, we investigated the cognitive effects of Zn^2+^ chelation in young mice. Previous studies have shown that intraperitoneal CQ administration promotes a robust reduction of the pool of chelatable Zn^2+^ in rodents ^33–38^. The main finding of the study is that a 2-week administration of CQ exerts adverse effects on learning and memory performances of treated mice. These results are in line with evidence supporting the critical role played by Zn^2+^ in the modulation of neuronal functioning ^38,45–47^. The study also indicates that the CQ-dependent effects are mediated by the inhibition of the BDNF neurotrophic axis.

### CQ administration affects BDNF signalling, synaptic plasticity, and behaviour

Zn^2+^ chelation profoundly affects mBDNF levels and the related signalling pathway. In CQ-treated animals, we observed a significant reduction in the total content of mBDNF. These changes were associated with a decrease of the total amount of the TrkB receptor. The decreased mBDNF levels that we observed were regionally restricted to the hippocampus, cortex, and striatum (Fig. 2). No effects were detected in the cerebellum, a finding that is in line with the hypothesis that the CQ-driven activity on BDNF signalling is selectively mediated by Zn^2+^ removal as the BDNF inhibition does not occur in a region, the cerebellum, that is devoid of pools of chelatable Zn^2+ 48^. These results also parallel the behavioural findings as the CQ-dependent cognitive effects are appreciated on a task, the MWM, that evaluates the hippocampus-related memory performances.

mBDNF exerts its neurotrophic action by binding to the TrkB receptor, and the study shows that CQ promotes a significant decrease in TrkB receptor levels. Unexpectedly, this result was not associated with a parallel reduction in the phosphorylated, activated, form of TrkB as we find that the pTrkB/TrkB ratio is unaffected by the drug treatment. The design of our experimental setting may help to explain the discrepancy. In our model, the molecular effects of CQ administration were assessed at the end of the 2-week treatment period. Within this time frame, it is conceivable that the CQ-mediated decrease in mBDNF is sufficient to induce a significant spine remodelling and/or elimination. This notion is, indirectly, supported by the observed reduction in PSD95 levels and further confirmed by the *in vitro* and *ex vivo* decreases of spine density that we observed (Fig. 3). Thus, the CQ-driven reduction of dendritic spines may have led to a decrease in the spine-associated transmembrane proteins including TrkB, thereby producing a net reduction of the receptor levels without affecting the pTrkB/TrkB ratio. The idea of defective TrkB signalling is further corroborated by the observed CQ-driven decrease in pERK5 levels, a kinase that acts downstream of the BDNF-TrkB neurotrophic axis and is known to promote neurogenesis and neuronal survival ^49,50^ but is also implicated in learning- and memory-related processes ^51,52^.

Our findings support a role for CQ-driven Zn^2+^ deprivation in the modulation of synaptic plasticity and dendritic spine pruning. CQ-treated hippocampal neurons show a reduced number of spines when compared with vehicle-treated sister cultures (Fig. 3). Our data are in line with the notion that mBDNF participates in structural plasticity by modulating the spine pruning and morphology ^53–55^. These plasticity-related phenomena represent the biophysical underpinning of the processes controlling the experience-dependent information storage occurring within the brain circuitry. Thus, the CQ-driven reduction of mBDNF levels may serve as one of the molecular mechanisms underlying the behavioural impairments observed in our model.

In the study, we investigated the Zn^2+^-dependent MMP activity with the aim of providing a mechanistic link between CQ-administration, Zn^2+^ chelation, and the reduction of mBDNF levels. Among MMPs, MMP-9 plays a permissive role in favouring structural plasticity and promote the dendritic spine expansion ^56^. Experimental evidence indicates that LTP-induced increase in spine size requires MMPs proteolytic activity ^22^ and, conversely, inhibition of MMPs has been shown to impair spatial learning in adult animals ^57,58^. In addition, recent findings have shown that MMP-9 participates, in a Zn^2+^ dependent manner, to BDNF maturation ^8,9^. In agreement with this notion, our *in gel* zymography experiments showed that CQ concentrations as low as 10 μM are sufficient to abolish the MMP gelatinolytic activity (Fig. 4). A further mechanistic link relies on the Zn^2+^-dependent transactivation of TrkB ^30^, an event that is likely occluded by CQ administration. While the mechanism is possible from a theoretical standpoint, it is unlikely to have occurred in our setting as we did not observe significant changes in TrkB phosphorylation and activation. In addition, a recent report has indicated that the receptor transactivation by Zn^2+^ occurs in in *vitro* settings and only marginally *in vivo* ^44^.

This Zn^2+^-dependent modulation of the MMP activity is of particular relevance when considering the cation role in synaptic plasticity. Zn^2+^ is particularly enriched within synaptic vesicles of a subset of hippocampal glutamatergic terminals ^27,47,59^. At these synapses, Zn^2+^ is released upon neuronal activity and plays a crucial role in the modulation of physiological and pathological processes ^28^. Substantiating this critical activity on plasticity and cognition, transgenic mice devoid of the synaptic zinc transporter-3 (ZnT-3 KO mice), and therefore unable to release Zn^2+^, show behavioural and molecular deficits that closely resemble the ones observed in our CQ-treated mice. Further suggesting a commonality of activities between transgenically- or pharmacologically-driven Zn^2+^ deprivation, ZnT-3 KO mice, like CQ-treated animals, show alterations in the BDNF-related pathways and plasticity markers ^60^.

In summary, CQ administration, by chelating Zn^2+^, reduced the MMP activity that, in turn, decreased BDNF maturation and defective structural plasticity, thereby leading to impaired cognitive performances.

### CQ administration modulates neurotoxic pathways in a pro-BDNF-independent manner

mBDNF is strongly related to its precursor form pro-BDNF. The decrease of mBDNF that we found was associated with the overexpression of the p75NTR receptor as well as with increased ERK_1,2_ signalling, a pathway that is activated by the pro-BDNF. Our WB data indicate that the treatment also produced decreased of pro-BDNF levels (Fig. 2G). These unexpected results prompted us to investigate whether CQ administration has affected the *BDNF* gene expression as previous studies have shown that changes in Zn^2+^ availability impinge on *BDNF* expression ^31,43^. qRT-PCR experiments, aiming at investigating the issue, indicated that the drug treatment did not alter *BDNF* mRNA levels. The additional targets of MMPs offer a possible explanation for these results. In addition to BDNF, MMPs are known to promote the cleavage and maturation of other pro-neurotrophins, including pro-NGF, a process that activates a deleterious p75NTR-related signalling ^61^. An additional, untested, possibility revolves around CQ effects on other intracellular Zn^2+^-dependent proteases that are involved in the BDNF maturation ^9,62^. Moreover, the reduction of mature neurotrophins, including mBDNF, can lead to a feed-forward increase of p75NTR expression and activation ^63^. Given the fact that TrkB and p75NTR mediate opposite effects ^11,64^, it is therefore conceivable that the mBDNF reduction unleashes a damaging increase in p75NTR levels (Fig. 4H). The hypothesis is unlikely in our setting as the enhanced p75NTR levels did not promote the activation of death pathways. The treatment did not result in increased neuronal toxicity, and the compound was found to be safe and well tolerated (Figs. 1B, and Supplementary fig. 2). Our p75NTR results, instead fit with previous findings showing that p75NTR-related signalling facilitates the activation of NMDAR-dependent long-term depression ^65,66^, negatively modulates dendritic morphology and spine density in hippocampal pyramidal neurons ^4,67^, and contributes to synaptic and memory deficits associated with neurodegenerative conditions ^68^. Similarly, aberrant activation of ERK_1,2_ not only results in the activation of death-related pathways but has been recently implicated in memory impairment in a mouse model of autism ^69^.

## Conclusions

Our findings support the notion that Zn^2+^, through the activation of the BDNF signalling cascade, is an important modulator of neuronal functioning and cognitive processes.

CQ has been employed in preclinical models of neurodegenerative conditions (AD and Huntington’s disease) with encouraging results ^70–73^. These robust preclinical findings were matched by positive effects in clinical trials exploring the use of CQ or other metal-protein-attenuating compounds (MPACs) in AD patients ^74–78^. Our study underscores the importance of choosing “when”, “where”, and “how much” modulation has to be applied when dealing with brain Zn^2+^ levels. Our findings do not challenge the importance of targeting Zn^2+^, with CQ or similar MPACs, as a therapeutic approach against neurodegenerative conditions but at the same time highlights the potential drawbacks of reducing brain Zn^2+^ availability in the early stages of development.

## Materials and Methods

### Animal model and treatment paradigm

All the procedures involving the animal handling and their care were approved by the University G. d’Annunzio Institutional Ethics Committee (CEISA protocol no. AD-301). *In vivo* procedures were performed in compliance with institutional guidelines and in accordance with national and international laws and policies. Mice were bred in the CeSI-MeT animal facility, single-housed, maintained on a 12 h light/dark cycle, and provided with*, ad libitum*, access to food and water. Behavioral tests were all performed during the early phases of light cycle (8 am – 11 am). C57Bl6 male mice were enrolled at 2.5 months of age (m.o.a.) and randomly assigned to a 2-week CQ or vehicle (see below) administration. Animals were treated daily with an intraperitoneal injection of CQ (30 mg/kg body weight) or vehicle. CQ was prepared by first dissolving the drug in DMSO (80 mg/ml) and then diluted in castor oil to a final concentration of 4 mg/ml (with a final 5% DMSO content).

Mice body weight was measured daily as a gross index of the overall safety of the treatment. A cohort of animals was employed for behavioural assessment, a second group which did not perform a behavioural evaluation, was employed for biochemical analysis.

### Neuronal cultures

Primary hippocampal neuronal cultures were prepared as previously described ^52,79^. Briefly, hippocampi were obtained from C57Bl6 fetuses at 15 or 16 embryonic days (E15 - E16). Alternatively, some experiments were performed on hippocampi obtained from CD1 mice. Hippocampi were dissected on ice and transferred in a 0.25% trypsin/EDTA solution for 10 min at 37 °C, then pelleted by centrifugation, and dissociated with a fire-polished glass pipette. The neuronal suspension was then diluted in Neurobasal medium supplemented with 0.5 mM l-glutamine, 5% fetal bovine serum, 5% horse serum, 1 × B27 and 0.2% penicillin/streptomycin and plated onto pre-treated laminin/poly-dl-lysine coated tissue culture glass coverslips. Three days after plating, non-neuronal cell growth was inhibited by adding 5 μM of cytosine arabinofuranoside, to obtain near-pure hippocampal cultures. 25% of the medium was replaced with fresh Neurobasal two times per week. Experiments were performed on cultures between 12 and 18 days in vitro (12 - 18 DIV).

### Morris Water Maze

The Morris Water Maze (MWM) test was performed as previously described ^52,80,81^. In brief, the MWM test was performed with a circular pool (Panlab; 1.2m diameter) filled with warm (22±1° C) water. The pool is placed in a room with several intra- and extra-maze visual cues for mice orientation. Mice were habituated to the testing room and placed on the platform 10 s before the first training session to reduce task-related stress. Mice underwent four trials per day for 4 consecutive days. 1.5 and 24 h after the end of the last training trial mice were tested for spatial memory performances. 1.5 and 24 h probe tests consisted of a 60 s free swim in the pool without the submerged platform. Probe sessions were digitally recorded and analyzed. The following parameters were measured: time to reach the location where the platform used to be (escape latency), number of crosses over the platform location (crosses), time spent in the target (T-target) or opposite (T-opposite) quadrants, average speed (cm/s), and total distance travelled during the LTM session.

### Tissue collection and reagents employed

Mice were euthanized after the 2-week treatment and tissue samples collected for further analysis. Brains were halved into two hemispheres, snap frozen by liquid nitrogen, and kept at −80 °C un l used. For Western blot (WB) analysis, each hemisphere was dissected into sub-regions (hippocampus, cortex, striatum, and cerebellum), snap frozen in liquid nitrogen, and stored at −80 °C un l sampling. For quantitative Real-Time PCR (qRT-PCR) hippocampi obtained from CQ- and vehicle-treated mice were stored in RNA*later* solution (ThermoFisher Scientific) until RNA extraction. For Golgi staining hemispheres were rinsed in PBS and immediately transferred to Golgi-Cox solution following manufacturer instructions (see below). All standard chemicals and reagents were, unless otherwise specified, purchased from Sigma-Aldrich.

### TSQ staining

TSQ staining was performed as described by Frederickson et al. with slight modifications ^82^. Briefly, 3 h after vehicle or CQ administration, mice were sacrificed, brains extracted, and cryosectioned without fixation. 6 μm thick slices were immediately stained with TSQ (100 μM; Santa Cruz Biotechnology) diluted in borate buffer [BB; containing (in mM): 100 boric acid, 25 sodium tetraborate, 75 NaCl, pH 9.5] for 90 s. Slices were then rinsed in PBS and put on coverslips. Images were acquired with a LSM 800 confocal microscope (Zeiss) by exciting the specimen with a 405 nm LED laser source. DRAQ5 (5 μM) was employed to counterstain the neuronal nuclei.

### Protein assay

Protein concentration was assayed by employing the Pierce BCA Protein Assay kit (Thermo scientific). Briefly, the assay is a detergent-compatible formulation based on bicinchoninic acid (BCA) for the colorimetric detection and quantitation of total protein. The method combines the reduction of Cu^2+^ to Cu^+^ ions by protein in alkaline medium (the biuret reaction) with the high sensitive and selective colorimetric detection of the cuprous cation by using a reagent containing BCA. The purple-colored reaction product of this assay is formed by the chelation of two molecules of BCA with one cuprous ion. This complex exhibits a strong absorbance at 562 nm.

### Western blot analysis

Brain regions were lysated in ice-cold RIPA buffer [pH 7.4 containing (in %): 0.5 sodium deoxycholate, 1 NP-40, 0.1 SDS, 1 protease and phosphatase inhibitor cocktails, and 5mM EDTA]. Protein lysates (10 μg) were separated by electrophoresis on 9-13% SDS–polyacrylamide gel and blotted onto polyvinyl difluoride (PVDF) membrane. Subsequently, membranes were blocked with 5% non-fat dry milk (Bio-Rad Laboratories) in Tris-buffered saline [TBS; containing (in mM): 20 Tris–HCl, 150 NaCl, pH 7.4] for at least 30 min at room temperature (RT). Membranes were incubated overnight with primary antibodies diluted in TBS containing 0,1% Tween 20 (TBS-T) and 5% non-fat dry milk.

The antibodies and their final concentration used were: anti-pro-BDNF 1:500 (Millipore), rabbit anti-BDNF 1:1000 (Abcam), rabbit anti-TrkB 1:1000 (Abcam), anti-pTrkB (Abcam), rabbit anti-PSD95 1:1000 (Cell Signaling), mouse anti-pERK_1,2_ 1:500 (Santa Cruz), rabbit anti-p75NTR 1:20000 (Abcam), rabbit anti-pERK5 1:1000 (Millipore), and rabbit anti-actin 1:2000 (Sigma). Specie-specific HRP conjugated IgG (1:10000; Vector Laboratories) were employed as secondary antibodies. The obtained chemiluminescent signals were visualized by ECL SuperSignal West Pico PLUS (Thermo scientific) and normalized to actin by using ImageJ software. Values are reported as relative units (R.U.) or phosphoprotein/total protein ratio calculated as: (phosphoprotein/loading control)/(total protein/loading control).

### Golgi staining

Hemispheres from vehicle- and CQ-treated mice were stained with an enhanced Golgi-Cox staining system (superGolgi Kit, Bioenno Tech), following manufacturer instructions. Briefly, 150 μm thick brain sections were cut using a vibratome (Leica) and mounted on adhesive microscope slides. Air-dried sections were stained, cleared in xylene, coverslipped in Eukitt mounting medium, and stored in the dark until analyzed. Images from secondary dendrites of CA1 pyramidal neurons were acquired with a DMRD microscope (Leica) equipped with a DFC550 CCD camera (Leica) and a 100x oil-immersion objective (1.3 NA; Leica). A trained experimenter blind to treatment manually counted the spines.

### Gelatin zymography

Gelatin zymography was performed as previously described ^32^. SDS-PAGE gels (7.5%) were prepared by incorporating 0.1% gelatin during acrylamide/bis-acrylamide polymerization. In all lanes, 10 μl of the supernatant, obtained from cultured hippocampi, was loaded. Gels were subjected to electrophoresis at 125 V constant. After electrophoresis, SDS was removed by washing gels for 30 min with a renaturating solution containing (in mM): 50 Tris-HCl, pH 7.4, and 2.5% Triton X-100. Gels were further washed with large volumes of ddH_2_O to remove residual Triton X-100. To assess MMPs activity gels were incubated overnight at 37° C in an activation buffer containing (in mM): 50 Tris-HCI, pH 7.4, 120 NaCl, 5 CaCl_2_. Accordingly to the experimental setting, activation buffer was supplemented with CQ (10 μM), TPEN (100 μM), or vehicle (DMSO 1%). Gels were then incubated for 30 min with a developing solution containing (in %): 0.1 Coomassie blue, 40 methanol, 10 acetic acid, 50 ddH_2_O. Staining was stopped when clear gelatinolytic bands appeared. Gels were photographed with a Gel-Doc apparatus (Biorad) and bands quantified by densitometry by using the Fiji distribution of ImageJ (v 1.51) ^83^.

### RNA extraction and qRT-PCR

The NucleoSpin Tissue RNA isolation kit (Macherey-Nagel) was employed for total RNA extraction. One μg of RNA was retro-transcribed using the High Capacity RNA-to-cDNA Kit (ThermoFisher Scientific). qRT-PCR was performed in a total volume of 25 μl containing 2X Maxima SYBR Green/ROX qPCR Master Mix (ThermoFisher Scientific), 1 μL of cDNA and 0.3 μM of each primer on an Abi 7900HT Sequencing Detection System (ThermoFisher Scientific). *BDNF* relative expression was corrected against GAPDH and HPRT1 that were used as endogenous controls. Amplification conditions were: 2 minutes at 50°, 10 minutes at 95°C, followed by 40 cycles of 15 seconds at 95°C and 1 minute at 60°C. A melting curve was run to assess the specificity of primers employed. Samples were run in triplicate. The genes relative fold changes were calculated by the ΔΔCt method.

The specific primers employed were: *BDNF* (F 5’-GAGACAAGAACACAGGAGGAAA-3’, R 5’-GACTAGGGAAATGGGCTTAACA-3’), *GAPDH* (F 5’-AACAGCAACTCCCACTCTTC-3’, R 5’-GTGGTCCAGGGTTTCTTACTC-3’) and *HPRT1* (F 5’-GGCCAGACTTTGTTGGATTTG-3’, R 5’-CGCTCATCTTAGGCTTTGTATTTG-3’).

### Neuronal viability assay

To exclude a possible neurotoxic effect of the compound, we assessed cell viability *in vitro*. Primary hippocampal neurons were exposed for up to 3 days to CQ (10 μM) or vehicle (DMSO). CQ toxicity was evaluated by lactate dehydrogenase (LDH) efflux assay as previously described ^84^.

Neuronal viability was also indirectly evaluated in Golgi stained *ex vivo* brain slices. Briefly, 20x magnification images of hippocampal and cortical areas were acquired and the number of stained neurons per field was counted and averaged.

### In vitro analysis of spine morphology

Spine morphology analysis was performed by transfecting primary hippocampal neurons with an EGFP containing construct. Transfection was performed at 7 DIV with Lipofectamine 2000 (ThermoFisher) following manufacturer instructions. Transfected cultures were subsequently treated with CQ or vehicle for 3 days. Dendritic processes of CQ- or vehicle-treated transfected neurons were imaged with a LSM 510 META confocal microscope (Zeiss) by exciting the specimen with a 488 nm Argon laser. z-stack imaging series were acquired and stored for further analysis. 3-dimensional reconstruction was obtained from stacked images and analyzed by employing the Fiji 3D Object Counter function.

### Statistical analysis

No methods were employed to predetermine sample size. Statistical analysis was performed by two-tailed Student’s t-test. For gelatin zymography ANOVA following Bonferroni post-hoc test was employed. The level of significance was set at p<0.05. Data were expressed as mean ± SEM.

## Acknowledgments

The authors are in debt with Angela Maria Vergnano for the helpful discussions and the critical revision of the manuscript. The authors thank Carlo Corona for helping with the gelatin zymography as well as Annalisa Nespoli and Manuela Iezzi for helping with brain cryosectioning. SLS is supported by research grants from the Italian Department of Education (PRIN 2011; 2010M2JARJ_005) and the Italian Department of Health (RF-2013–02358785 and NET-2011-02346784-1).

## Author contributions

SLS and VF conceived and designed the experiments. MB, VF, and AG performed *in vivo* treatment and sample collection. MB performed behavioural testing. VF performed *in vitro* spine analysis. AG performed and analyzed gelatin zymography, TSQ staining and LDH assay. AG and NM performed and analyzed Golgi staining experiment. VC and AC performed and analyzed WB experiments. MdA and VG performed and analyzed qRT-PCR experiment. VF, AG, MP, MI, AM, SDP, and SLS analyzed and interpreted the data. VF, AG, and SLS wrote the manuscript. SLS supervised the study. All authors discussed, revised, and approved the final version of the manuscript.

## Data availability

The data of the study are available from the corresponding author upon reasonable request.

---

**Supplementary figure 1. CQ administration reduces chelatable zinc in the brain.** TSQ staining was employed to assess levels of chelatable Zn^2+^ in the hilus of the hippocampal dentate gyrus (DG). (A) Representative images of brain sections obtained from vehicle- (left) and CQ-treated (right) mice. (B) Magnification of the hilus of the DG as in (A). (B) The bar graph shows the quantification obtained from images in A (TSQ normalized fluorescence in vehicle-treated 1.00±0.11 vs 0.49±0.04 in CQ-treated mice, n=6-8 brain slices from vehicle- and CQ-treated mice, p<0.001). “***” indicates p<0.001

**Supplementary figure 2. CQ supplementation does not affect neuronal viability.** (A) Representative images of Golgi-stained brain sections obtained from vehicle- (left) and CQ-treated (right) mice. The Golgi-staining was employed as an indirect index of neuronal viability. The viability was evaluated as the average number of neurons per field in slices obtained from vehicle- and CQ-treated mice. (B-C) Bar graphs depict the quantification of the average number of Golgi-stained neurons per field (Hippocampus: 15.62±0.87 for control vs 15.72±1.55 for CQ-treated mice, n=4 per condition, p=0.87; Cortex: 19.20±1.92 for control vs 25.52±2.30 for CQ-treated mice, n=4 per condition, p=0.08). (D) Bar graph depicts the viability of cultured hippocampal neurons exposed to CQ (10 μM) or vehicle for 3 days. Neuronal viability was assessed by LDH efflux assay and results normalized to a set of experiments in which almost complete neuronal death was achieved by exposing cells to toxic levels of NMDA (300 μM + 10 μM glycine) for 24h (Control 30.94±2.88 vs 30.08±2.81 in CQ-treated cultures, n=9 per condition, p=0.83).

**Supplementary table 1.** Statistics for the WB experiments shown in main text.

|  |  | Control | Clioquinol |  |  |  |
| --- | --- | --- | --- | --- | --- | --- |
|  |  | Mean±SEM | Mean±SEM | p | n | Significance |
| **BDNF** | Hippocampus | 1.40±0.07 | 0.91±0.06 | 0.002 | 4 | ** |
|  | Cerebellum | 0.21±0.04 | 0.29±0.07 | 0.34 | 4 |  |
|  | Cortex | 1.81±0.06 | 1.30±0.14 | 0.015 | 4 | * |
|  | Striatum | 0.86±0.09 | 0.50±0.06 | 0.022 | 4 | * |
| **TrkB** | Hippocampus | 1.13±0.05 | 0.66±0.12 | 0.013 | 4 | * |
|  | Cerebellum | 0.54±0.06 | 0.49±0.06 | 0.57 | 4 |  |
|  | Cortex | 2.00±0.08 | 1.71±0.07 | 0.03 | 4 | * |
|  | Striatum | 1.32±0.07 | 0.78±0.04 | 0.001 | 4 | ** |
| **pTrkB/TrkB** | Hippocampus | 0.74±0.09 | 0.84±0.21 | 0.66 | 3 |  |
|  | Cerebellum | 0.38±0.04 | 0.54±0.19 | 0.47 | 3 |  |
|  | Cortex | 0.56±0.07 | 0.43±0.01 | 0.15 | 3 |  |
|  | Striatum | 0.49±0.01 | 0.55±0.09 | 0.56 | 3 |  |
| **PSD95** | Hippocampus | 0.89±0.03 | 0.69±0.04 | 0.02 | 3 | * |
|  | Cerebellum | 0.24±0.03 | 0.20±0.03 | 0.45 | 3 |  |
|  | Cortex | 1.62±0.04 | 1.29±0.04 | 0.007 | 3 | ** |
|  | Striatum | 1.20±0.05 | 0.71±0.08 | 0.008 | 3 | ** |
| **pERK5/ERK5** | Hippocampus | 0.91±0.08 | 0.34±0.03 | 0.002 | 3 | ** |
|  | Cerebellum | 0.41±0.03 | 0.43±0.01 | 0.57 | 3 |  |
|  | Cortex | 0.80±0.03 | 0.52±0.06 | 0.019 | 3 | * |
|  | Striatum | 0.54±0.08 | 0.26±0.02 | 0.03 | 3 | * |
| **proBDNF** | Hippocampus | 1.22±0.06 | 0.91±0.05 | 0.02 | 3 | * |
|  | Cerebellum | 0.28±0.04 | 0.20±0.02 | 0.18 | 3 |  |
|  | Cortex | 1.19±0.10 | 0.64±0.07 | 0.01 | 3 | * |
|  | Striatum | 0.63±0.05 | 0.23±0.05 | 0.006 | 3 | ** |
| **p75NTR** | Hippocampus | 1.16±0.07 | 1.49±0.04 | 0.01 | 3 | * |
|  | Cerebellum | 0.30±0.03 | 0.44±0.06 | 0.11 | 3 |  |
|  | Cortex | 2.47±0.03 | 2.74±0.02 | 0.002 | 3 | ** |
|  | Striatum | 0.22±0.03 | 0.93±0.03 | <0.001 | 3 | *** |
| **pERK<sub>1,2</sub>/ ERK<sub>1,2</sub>** | Hippocampus | 0.54±0.05 | 0.88±0.08 | 0.03 | 3 | * |
|  | Cerebellum | 0.20±0.01 | 0.24±0.01 | 0.10 | 3 |  |
|  | Cortex | 1.59±0.13 | 2.13±0.07 | 0.02 | 3 | ** |
|  | Striatum | 0.98±0.01 | 1.22±0.069 | 0.03 | 3 | ** |

Data are represented as mean ± SEM. “\*” indicates $p < 0.05$ , “\*\*” indicates $p < 0.01$ , and “\*\*\*” indicates $p < 0.001$ .

